# Expert Panel Curation of 31 Genes in Relation to Limb Girdle Muscular Dystrophy

**DOI:** 10.1101/2024.05.03.592369

**Authors:** Shruthi Mohan, Shannon McNulty, Courtney Thaxton, Marwa Elnagheeb, Emma Owens, May Flowers, Teagan Nunnery, Autumn Self, Brooke Palus, Svetlana Gorokhova, April Kennedy, Zhiyv Niu, Mridul Johari, Alassane Baneye Maiga, Kelly Macalalad, Amanda R. Clause, Jacques S. Beckmann, Lucas Bronicki, Sandra T. Cooper, Vijay S. Ganesh, Peter B. Kang, Akanchha Kesari, Monkol Lek, Jennifer Levy, Laura Rufibach, Marco Savarese, Melissa J. Spencer, Volker Straub, Giorgio Tasca, Conrad C. Weihl

**Affiliations:** Department of Genetics, University of North Carolina, Chapel Hill, NC, USA; Aix Marseille Univ, INSERM, MMG, U 1251, Marseille, France; Department of Medical Genetics, Timone Children’s Hospital, APHM, Marseille, France; The Hospital for Sick Children, Toronto, ON, Canada; Department of Laboratory Medicine and Pathology, Mayo Clinic; Harry Perkins Institute of Medical Research, Centre for Medical Research, University of Western Australia, Nedlands, WA, Australia; Folkhälsan Research Center, Department of Medical and Clinical Genetics, Medicum, University of Helsinki, Finland; University of Sciences, Techniques and Technologies of Bamako, Mali; Washington University School of Medicine in St. Louis; University of Lausanne, Lausanne, Switzerland; Children’s Hospital of Eastern Ontario, Ottawa, Ontario, Canada; Kids Neuroscience Centre, Children’s Hospital at Westmead; School of Medical Sciences, Faculty of Medicine and Health, The University of Sydney; Functional Neuromics, Children’s Medical Research Institute, Westmead, NSW, Australia; Center for Mendelian Genomics, Broad Institute of MIT and Harvard, Cambridge, MA; Department of Neurology, Brigham and Women’s Hospital, Boston, MA; Greg Marzolf Jr. Muscular Dystrophy Center and Department of Neurology, University of Minnesota, Minneapolis, MN, USA; Illumina Inc, San Diego, CA, USA; Department of Genetics, Yale University School of Medicine, New Haven, CT, USA; Coalition to Cure Calpain 3, Westport, CT, USA; Jain Foundation, Seattle, WA USA; David Geffen School of Medicine at UCLA, Los Angeles, CA, USA; John Walton Muscular Dystrophy Research Centre, Newcastle University and Newcastle Hospitals NHS Foundation Trusts, Newcastle Upon Tyne, UK

**Keywords:** limb girdle muscular dystrophy, LGMD, gene-disease validity, next-generation sequencing, ClinGen

## Abstract

**Introduction:** Limb girdle muscular dystrophies (LGMDs) are a group of genetically heterogeneous autosomal conditions with some degree of phenotypic homogeneity. LGMD is defined as having onset >2 years of age with progressive proximal weakness, elevated serum creatine kinase levels and dystrophic features on muscle biopsy. Advances in massively parallel sequencing have led to a surge in genes linked to LGMD.

**Methods:** The ClinGen Muscular Dystrophies and Myopathies gene curation expert panel (MDM GCEP, formerly Limb Girdle Muscular Dystrophy GCEP) convened to evaluate the strength of evidence supporting gene-disease relationships (GDR) using the ClinGen gene-disease clinical validity framework to evaluate 31 genes implicated in LGMD.

**Results:** The GDR was exclusively LGMD for 17 genes, whereas an additional 14 genes were related to a broader phenotype encompassing congenital weakness. Four genes (*CAPN3, COL6A1, COL6A2, COL6A3*) were split into two separate disease entities, based on each displaying both dominant and recessive inheritance patterns, resulting in curation of 35 GDRs. Of these, 30 (86%) were classified as Definitive, 4 (11%) as Moderate and 1 (3%) as Limited. Two genes, *POMGNT1* and *DAG1*, though definitively related to myopathy, currently have insufficient evidence to support a relationship specifically with LGMD.

**Conclusions:** The expert-reviewed assertions on the clinical validity of genes implicated in LGMDs form an invaluable resource for clinicians and molecular geneticists. We encourage the global neuromuscular community to publish case-level data that help clarify disputed or novel LGMD associations.

## Introduction

Limb girdle muscular dystrophies (LGMDs) comprise a group of disorders characterized by weakness and progressive wasting of proximal limb muscles ^1^. Although phenotypically similar with regard to pattern of weakness, LGMDs can show variable expressivity, ranging from severe, early onset to mild, late onset forms ^2^. Prior to advances in massively parallel sequencing (MPS) methods, LGMD diagnoses were based on linkage studies in large pedigrees, clinical phenotyping, muscle pathology findings and candidate gene screening via Sanger sequencing ^3^. More than 30 genes are currently identified as being implicated in LGMD, with both autosomal dominant (AD) and autosomal recessive (AR) inheritance patterns ^4^. LGMDs have been classified into subtypes based on the causative gene and inheritance pattern, and this classification system and nomenclature have undergone revisions over the years ^5^. Revised, current nomenclature defines AD (LGMD D1-5) and AR (LGMD R1-28) disease entities. Notably, the LGMD nomenclature includes some genes that are related to a broader phenotypic spectrum. For example, a group of myopathies associated with a reduction in α-dystroglycan glycosylation can range from congenital onset with early lethality to mild, adult-onset weakness. In these cases, the single disease entity for curation may be more appropriately defined as a phenotypic spectrum that encompasses LGMD rather than as pure LGMD. The rationale for ascribing a disease gene to an LGMD subtype has not been systematic and most recently was achieved by expert consensus. This recent effort to reclassify LGMDs removed 10 genes previously deemed LGMD and added an additional 5 genes ^5^.

With the advances in MPS technologies, the ability to identify genomic variants has increased tremendously. However, accurate interpretation of the detected genomic variants is crucial for molecular diagnostics. Moreover, accurate variant interpretation is not possible without understanding the clinical validity of the implicated GDR. To this end, the Clinical Genome Resource (ClinGen) ^6^, an NIH-funded initiative to build an authoritative central resource to define the clinical relevance of genes and variants for use in precision medicine and research, has developed a semi-quantitative framework to assign clinical validity to gene-disease relationships ^7^. The ClinGen Muscular Dystrophies and Myopathies gene curation expert panel (MDM GCEP, formerly Limb girdle muscular dystrophy GCEP) applied this framework to assess the strength of evidence associated with genes asserted as LGMD in the revised LGMD nomenclature. Gene-disease validation is an essential step towards generating targeted testing panels ^8^ and establishing variant pathogenicity.

## Methods

### Muscular dystrophies and myopathies gene curation expert panel (Formerly Limb girdle muscular dystrophy gene curation expert panel)

The MDM GCEP was convened initially as the LGMD GCEP in 2019 as part of the ClinGen Neurological (formerly Neuromuscular) Clinical Domain Working Group to curate genes asserted with an LGMD disease relationship and has now expanded its scope of work to curate genes causing myopathies and other muscular dystrophies. It is an international collaboration of members from 14 institutions across Australia, Canada, Finland, France, Italy, Mali, Switzerland, the United Kingdom and the United States. The MDM GCEP consisted of 8 biocurators, 1 coordinator, and 14 experts with clinical, molecular genetic and/or research backgrounds that have contributed significantly to the growth of the LGMD field. The MDM GCEP webpage can be accessed at https://clinicalgenome.org/affiliation/40151/.

### Gene list for curation

The MDM GCEP focused on evaluating 35 GDRs related to LGMD or a disease spectrum including LGMD (Table 1). We included 31 genes in our initial scope of work listed by the 229th European Neuromuscular Centre (ENMC) International workshop on LGMDs as being related to LGMD ^5^. This workshop proposed new nomenclature and classification for LGMD subtypes (LGMDD1-5 and LGMDR1-24 at the time of workshop report) and a consensus definition of LGMD. In addition, we assessed two genes linked to LGMD by the Online Mendelian Inheritance in Man database (OMIM), *BVES* (LGMDR25) and *POPDC3* (LGMDR26). Five other included genes (*COL6A1, COL6A2, COL6A3, LAMA2, TTN*) were curated for their GDR by the ClinGen Congenital Myopathies GCEP, with the scope of the MDM GCEP to assess whether LGMD formed a part of the disease spectrum.

### Definition of LGMD for the purpose of gene curation

LGMDs range from severe forms with onset in the first decade and rapid progression (resembling Duchenne muscular dystrophy) to milder forms with late onset and slower progression. Typically, LGMD is characterized by progressive weakness, predominantly of the proximal limb muscles. The initial presentations are typically weakness of the hip and proximal leg muscles. Affected individuals usually have normal early motor and intellectual milestones. Cardiac involvement in the form of dilated or hypertrophic cardiomyopathy and dysrhythmias can also be present. The facial muscles are typically spared or only minimally involved. Muscle enzymes such as creatine kinase are typically elevated in serum, often with a level greater than twice the upper limit of the normal range identified in controls. Muscle biopsies typically have dystrophic features, but can also show milder myopathich characteristics. Electrodiagnostic features typically include myopathic motor unit morphologies and recruitment patterns.

For the purposes of gene curation, we defined that the LGMD clinical criteria include: 1) progressive weakness in limb girdle and proximal musculature; 2) independent ambulation by the age of 2 years; 3) elevated serum CK levles of greater than twice the upper limit of the reference range; 4) muscle biopsy with dystrophic or myopathic features.

### MDM GCEP gene-disease-mode of inheritance curation process

The first step in the gene curation process was “precuration”, which entailed evaluating and identifying the most appropriate disease entity and inheritance pattern in relation to which gene is curated. The MDM GCEP applied the ClinGen lumping and splitting guidelines ^9^ to assess genes reported to be related to more than one disease entity in OMIM or the literature. We took into consideration the multiple assertions, phenotypic variability, inheritance patterns and molecular mechanism(s) for pathology to inform whether multiple disease assertions should be lumped together for a gene curation or split to be curated separately. Upon performing this analysis, *CAPN3, COL6A1, COL6A2*, and *COL6A3* were split into two separate disease entities each, based on both autosomal dominant and recessive inheritance patterns and all other genes were curated for a single disease entity.

The MDM GCEP determined the clinical validity for GDRs using the ClinGen Gene Curation Standard Operating Procedure, Version 7-9 (SOP versions available at https://clinicalgenome.org/curation-activities/gene-disease-validity/training-materials/), based on the framework described in Strande & Riggs et al. ^7^. This framework comprises a semi-quantitative scoring system that uses an evidence-based approach to classify GDRs based on the strength of evidence supporting the causative role of the gene. Each evidence piece is awarded a certain number of points, which cumulatively yield overall classifications as follows: Definitive (12-18 points, with replication over time), Strong (12-18 points), Moderate (7-11 points), and Limited (1-6 points). While Definitive and Strong classifications require the same overall score, the Definitive classification also requires that the role of the gene in the disease has been repeatedly demonstrated in research and clinical diagnostic settings and upheld over time (a minimum of three years). In the absence of evidence of the gene’s role in human disease, the category of “No known disease relationship” can be applied, while the Disputed and Refuted classifications can be applied for GDRs with evidence contradicting the gene’s role in disease. In addition to evaluating GDRs, the MDM GCEP also assessed whether LGMD was a specific presentation within the disease spectrum for a particular gene and defined the threshold as four independent cases meeting the typical LGMD clinical criteria listed previously.

The ClinGen framework recommends that curations reaching a Limited through Strong classification be re-curated every two to three years to evaluate new evidence that could potentially upgrade or downgrade the existing classification for that curation. In addition, the reconvening of the GCEP for re-curation purposes also provides the opportunity for the group to identify additional and/or newly discovered genes to curate in relation to LGMD.

A systematic assessment of the available genetic and experimental evidence from published literature for every GDR was recorded in the ClinGen gene curation interface (GCI) by biocurators, leading to a provisional classification. The curated genetic evidence included variants in the gene of interest identified in patients with the disease entity being curated for, which included case reports, family studies and linkage analyses. The experimental category incorporated functional evidence that supported the role of the gene in the disease of interest, including biochemical function, protein interactions, expression, functional alterations in patient or non-patient cells, and animal models that recapitulated the disease and/or showed rescue of phenotype with the introduction of wild-type gene product. A maximum of 12 points were assigned for genetic evidence and 6 points for experimental evidence for a combined maximum of 18 points. Subsequently, the curated evidence was reviewed and discussed on GCEP conference calls, and once a consensus was reached among the experts, the curation was approved and published directly to the ClinGen website (https://www.clinicalgenome.org/affiliation/40151/). The non-Definitive classifications are likely to change with time as re-curation efforts are undertaken and, therefore, up-to-date curation data and classifications can be viewed by searching by the gene name on clinicalgenome.org.

### Collaboration with the ClinGen Congenital Myopathies GCEP

Of the 31 genes identified as high priority for curation by the MDM GCEP, five were also in the scope of work of the ClinGen Congenital Myopathies GCEP: *COL6A1, COL6A2, COL6A3, LAMA2*, and *TTN*. Following curation of these genes by the Congenital Myopathies GCEP, the MDM GCEP reviewed the lumping and splitting assessment, genetic and experimental evidence, and overall gene-disease classification proposed by the Congenital Myopathies GCEP for each GDR and recommended edits or additions to the curations to represent the perspectives of the MDM GCEP. Modifications to the curations were reviewed and approved by the Congenital Myopathies GCEP and, in the case of the *TTN* curation, a collaborative meeting with representatives from both panels was convened to finalize the curation.

### Mondo disease ontology selection and restructure of terms related to LGMD phenotypes

Each gene curation was related to a disease entity selected from the Monarch Disease Ontology (Mondo) ^10^ in alignment with the lumping and splitting determination. In the event that the MDM GCEP identified a need for the addition of a new term to Mondo to represent the selected disease entity for curation, requests for new Mondo terms were submitted to the Monarch Initiative’s Mondo GitHub (https://github.com/monarch-initiative/mondo), which allows for review and input from the Mondo user community.

## Results

The ClinGen MDM GCEP evaluated 31 genes implicated in LGMD, leading to 35 GDRs. Of these GDRs, 30 (86%) were classified Definitive, 4 (11%) were classified Moderate and 1 (3%) was classified Limited (Figure 1A). No gene curations reached a Strong, No Known Disease Relationship, Disputed or Refuted classification. Twenty-six genes were curated for a single disease entity, including 14 genes for which the curated disease was a lumped entity, while four genes were split into separate curations for different disease entities, based on the inheritance pattern (see Table 1).

**FIGURE 1.**
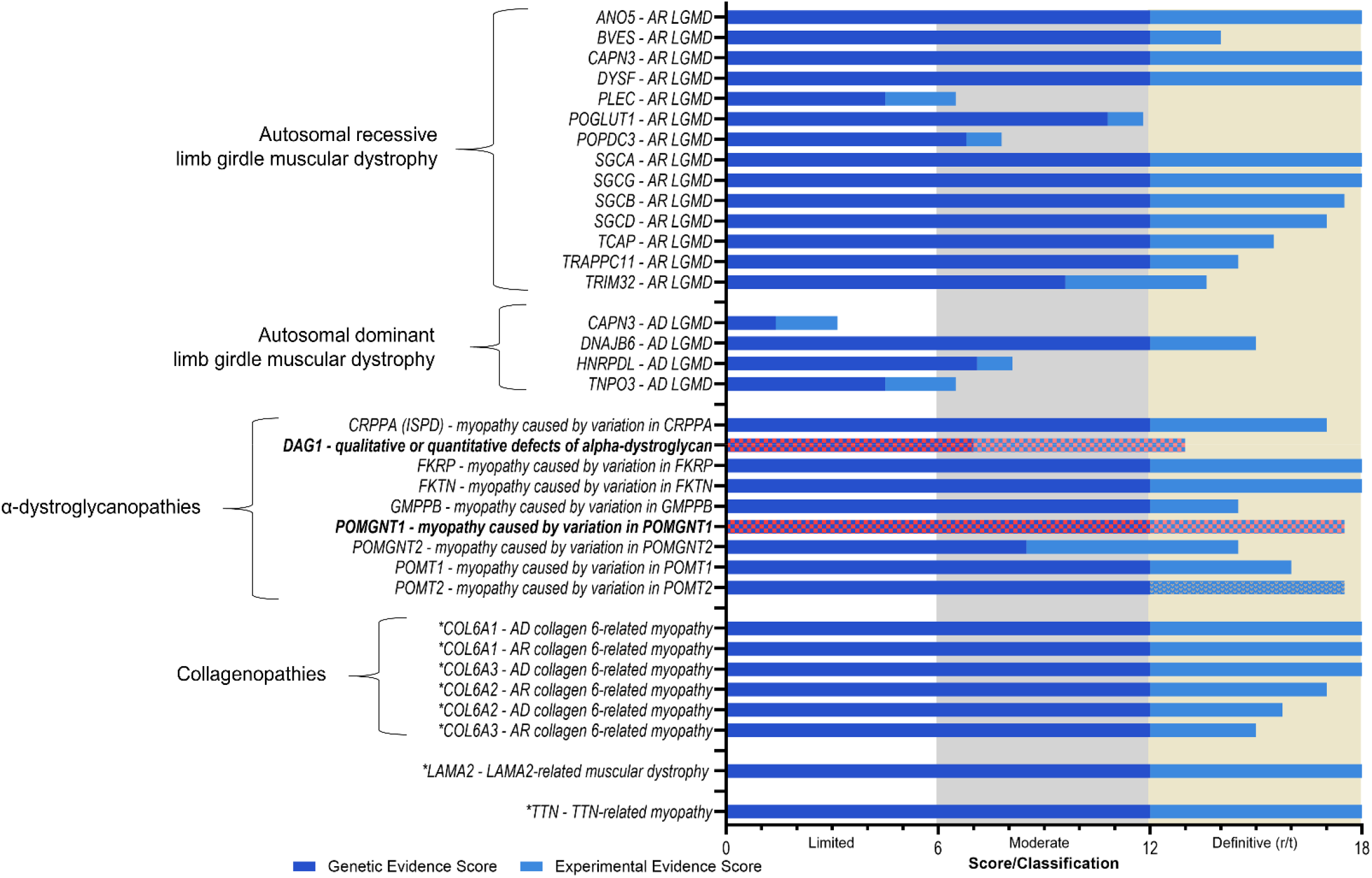
Curation classifications of the 35 gene-disease relationships curated by the MDM GCEP. Genetic and experimental evidence scores of the 35 gene-disease relationships. *POMGNT1* and *DAG1*, though definitively related to myopathy, currently have insufficient evidence to support a relationship specifically with LGMD and are noted in red. *denote genes co-curated by the congenital myopathy and MDM GCEP.

Eighty-six percent of LGMD GDRs reached a Definitive classification, with a majority (25/30) receiving the maximum 12 points for the genetic evidence category (Figure 1A and Table 1). Scores for experimental evidence ranged from 1 – 6 points, and 14/30 Definitive curations obtained all 6 points for the experimental category. Twelve Definitive curations received maximum scores in both genetic and experimental categories.

Among the four curations with Moderate classifications, though genetic evidence scores ranged between 4.5 - 7.1 points, experimental evidence was scarce, amounting to 1 – 2 points. In the case of the GDR for AD-*CAPN3* (LGMDD4) that had a Limited classification, both genetic and experimental evidence were limited, accumulating less than 2 points for genetic and experimental evidence (see Table 1).

The MDM GCEP re-curated four GDRs that reached a Limited or Moderate classification when they were originally curated at least two years prior. Curation of additional published evidence in the time elapsed, and rescoring of evidence based on the latest version (SOP v9) of the ClinGen gene-disease validity framework, led to the upgrading of the *DAG1* curation from Moderate to Definitive and of the *TNPO3* and *POPDC3* curations from Limited to Moderate classifications. There was no change to the classification of *PLEC* (Moderate) despite an increase in the genetic evidence scores.

Five genes overlapping Congenital Myopathies and MDM GCEPs (*COL6A1, COL6A2, COL6A3, LAMA2*, and *TTN)*, were linked to a disease spectrum that included both myopathy and LGMD phenotypes and thus were curated by the two panels in collaboration. The MDM GCEP reviewed preliminary curation data provided by the Congenital Myopathy GCEP and offered suggested modification to the lumping and splitting assessments associated with *COL6* and *TTN* genes. The MDM GCEP provided evidence for *LAMA2* and *TTN* gene curation related to published cases meeting the MDM GCEP phenotype definition of LGMD. Thus, the MDM GCEP is recognized as a secondary contributor to all eight GDR curations.

## Discussion

Using the ClinGen gene-disease validity framework, the MDM GCEP (formerly LGMD GCEP) classified 35 GDRs for 31 genes asserted in the ENMC revised LGMD nomenclature, providing the community with quality assessments of the strength of evidence supporting a gene-disease relationship and LGMD as a specific presentation within the disease spectrum. For genes that were asserted to be involved in only a specific LGMD subtype (e.g. LGMDR1), the MDM GCEP chose to curate for the disease entity of “Limb-Girdle Muscular Dystrophy”, and depending on the inheritance pattern, included “autosomal recessive” or “autosomal dominant” as a prefix. For genes implicated in more than a single disease entity, including phenotypes on a spectrum beyond LGMD, more inclusive nomenclature was used to indicate the lumped disease entity, also incorporating the gene name to be more specific and consistent with the recommendations from the ClinGen disease naming advisory committee, e.g., collagen 6-related myopathy, myopathy caused by variation in *POMT1*.

Four genes (*CAPN3, COL6A1, COL6A2, COL6A3*) were split based on differing inheritance patterns and curated as two separate GDRs for each gene. For each of these genes, the pathogenic mechanism of AR versus AD was deemed distinct. The mechanism for *CAPN3, COL6A* AR disease is a loss of function relating to a loss of protein expression or function. However in the case of AD inheritance, the mechanism is dominant negative for the *COL6A*, resulting in impaired collagen assembly ^11^. Although less clear, the dominant *CAPN3* mutations may also behave in dominant-negative manner resulting in decreased proteolytic activity or stability of the wild-type calpain-3 protein ^12^.

A primary goal of the MDM GCEP was to specify phenotypic criteria for assertion of an LGMD phenotype. For curation purposes, we defined LGMD phenotypic criteria as: 1) progressive weakness in limb musculature (pelvic and shoulder girdle); 2) independent ambulation by the age 2 years; 3) elevated CK of >2 times the upper limit of normal; 4) muscle biopsy with dystrophic or myopathic features. It is important to emphasize that these phenotypic criteria are not an attempt to define the phenotypic spectrum of LGMD disease genes, which often are part of a broader phenotypic spectrum that overlaps to varying degrees with congenital and distal myopathies and other phenotypic patterns of weakness. Moreover, the criteria were not intended to supplant diagnositic approaches that may include other modalities utilized in clinical practice such as electrodiagnostic studies or muscle magnetic resonance imaging. Rather, these criteria were refined by the MDM GCEP to provide a pragmatic, minimal list of core phenotypic criteria necessary to assert a gene-LGMD relationship.

Of the GDRs evaluated for an LGMD disease entity, extensive curation evidence was available for 12 Definitive GDRs (Figure 1B), which allowed scoring the maximum scores for genetic and experimental evidence. *CAPN3*, which has been reported in the autosomal dominant (AD) inheritance pattern in a few multi-generational pedigrees, was assessed as split curations for LGMD inherited in AD and AR patterns. The AR curation reached a Definitive classification, while the AD curation ended up being Limited due to the paucity of families identified and an unclear pathogenic mechanism.

Variation in multiple genes encoding proteins involved in the addition of O-linked oligosaccharides to α-dystroglycan (*CRPPA, FKRP, FKTN, GMPPB, POMGNT1, POMGNT2, POMT1*, and *POMT2)* have been implicated in muscular dystrophy, with the spectrum of phenotypes ranging from a severe muscle disorder also linked to brain and eye anomalies and, in some cases, intellectual disability, to a milder isolated LGMD presentation. The MDM GCEP recommended changes to MONDO classifications in order to reflect this spectrum (Figure 2). Each of these genes was curated by the MDM GCEP for their relationship with myopathies and reached a definitive classification. Importantly, although the LGMD phenotype has been asserted in the literature for variants in the *POMGNT1* gene, the reported individual did not meet the LGMD phenotype definition set forth by the GCEP ^13^ or the reported variants were considered unlikely to be causative ^14, 15^. *DAG1* was curated for the entity “qualitative or quantitative defects of α-dystroglycan”. This curation reached a Definitive classification upon recuration in March 2023. However, there were <4 probands in the curated literature meeting the LGMD phenotype criteria ^16-18^, and similar to *POMGNT1, DAG1* has limited evidence to support a causative relationship with an LGMD phenotype. It is important to note that the availability of more convincing evidence in the future could change this relationship. The MDM GCEP urges the community to publish more cases meeting the LGMD phenotype carrying causative variants in these genes.

**FIGURE 2.**
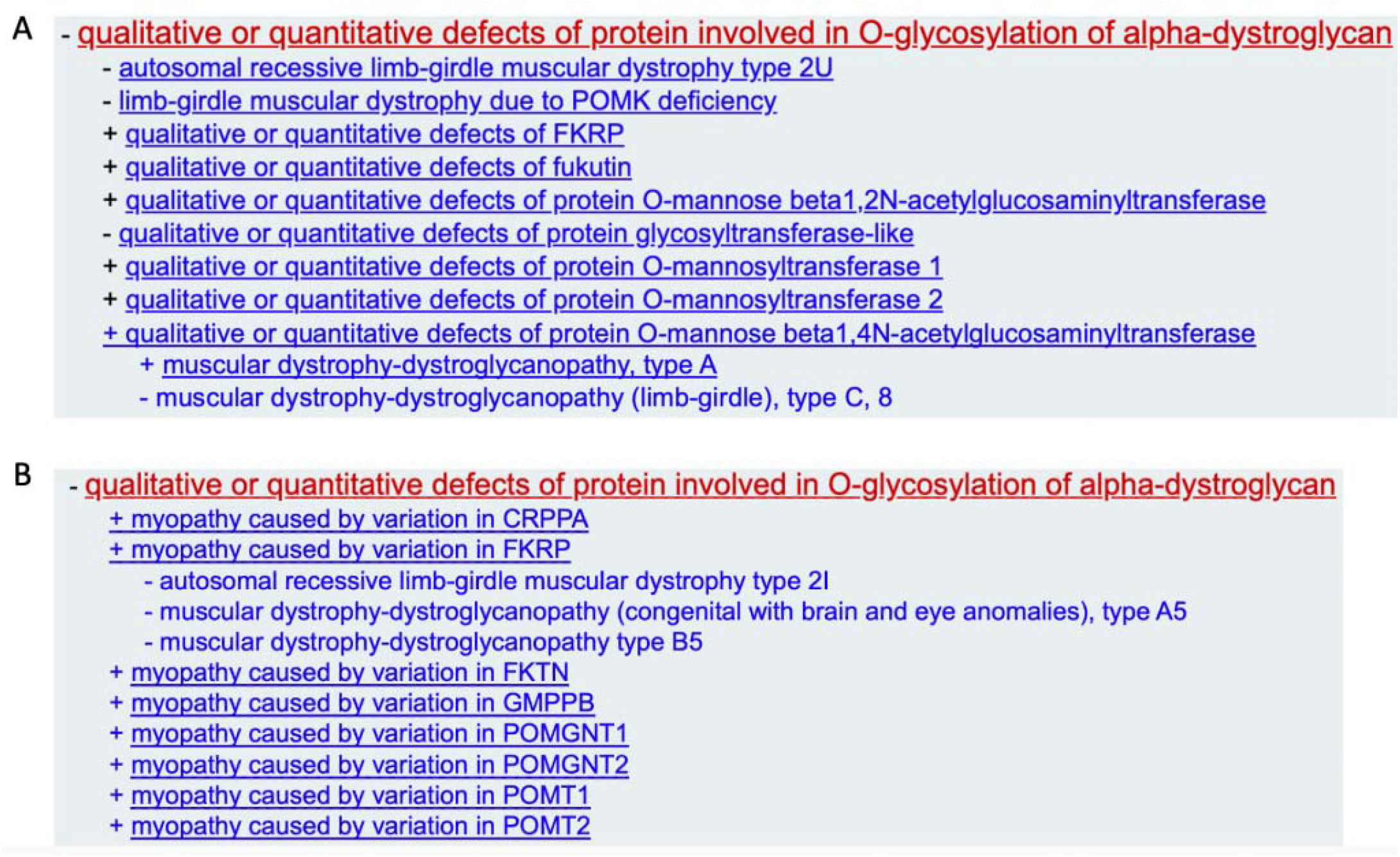
Mondo revision recommended by the MDM GCEP for the muscular dystrophy-dystroglycanopathy genes. A) Previous parent (+) and child (-) terms for α-dystroglycanopathy genes. B) Proposed parent and child terms for α-dystroglycanopathy genes.

AR LGMD due to *PLEC* variation is typically characterized by early childhood onset of proximal muscle weakness and atrophy, notably without skin involvement ^19^. In addition to an isolated LGMD phenotype, *PLEC* has been reported in relation to epidermolysis bullosa simplex with muscular dystrophy (EBS5B). The MDM GCEP distinguished the molecular mechanism of EBS5B as primarily nonsense, out-of-frame insertions or deletions within exon 31 and 32, leading to premature protein termination of all plectin isoforms. In contrast, the underlying mechanism for isolated LGMD without skin involvement appears to be recessive truncating variants specifically in exon 1f that encodes the plectin 1f isoform that is only expressed in skeletal muscle. A homozygous founder variant, a 9-base pair deletion, has been reported in all the probands scored for this curation ^19, 20^. Additional variants in *PLEC* have been noted in the literature in patients with asserted LGMD ^21-24^ however, the curation did not include these probands since they harbored missense variants or variants in other transcripts that did not solely impact the plectin 1f isoform and were considered to have an EBS syndrome in addition to muscle weakness.

*HNRNPDL* reached a Moderate classification in relation to AD LGMD, and all probands and families scored for genetic evidence were heterozygous for variants at the same residue, Asp378 ^25^. Additional caution is warranted when interpreting variants at other positions in this gene. *TNPO3* was recurated in March 2023, and while this curation was initially classified as Limited, it was upgraded to Moderate upon recuration. Evidence that aided in this upgrade came from a Drosophila model published in 2021 ^26^, which showed partial recapitulation of the impaired muscle function typical of patients with AD LGMD due to *TNPO3* variation.

The collaboration between the MDM GCEP and the Congenital Myopathies GCEP on five genes (*COL6A1, COL6A2, COL6A3, LAMA2*, and *TTN*) led to the MDM GCEP being a secondary contributor on the eight Definitive curations. Genetic analysis in patients with collagen 6-related myopathies revealed both dominant and recessive defects in one of the *COL6A1, COL6A2*, or *COL6A3* genes that affect the final Collagen VI protein. On the basis of clinical as well as genetic findings, Ullrich congenital muscular dystrophy, Bethlem myopathy and LGMD can justifiably be regarded as a continuous spectrum of disease ranging from congenital to late adolescent/early adulthood onset rather than distinct clinical entities ^27^. For each of these curations, the MDM GCEP identified at least 4 independent probands that supported a phenotypic presentation consistent with LGMD.

In conclusion, the majority of GDRs reached a Definitive classification upon assessment by the MDM GCEP. These expert-reviewed assertions on the clinical validity of genes implicated in LGMDs form an invaluable resource for clinicians and laboratory diagnosticians. The Moderate and Limited relationships will be reevaluated by the GCEP in 2-3 years’ time, along with assessing any additional and/or novel genes reported in relation to LGMD. Indeed, since the inception of the MDM GCEP, three new LGMD genes have been published and assigned LGMDR27-29 (*JAG2, HMGCR* and *SNUPN* respectively) by OMIM ^28-31^. Systematic evaluation of GDRs related to LGMD is necessary to standardize the genes included on genetic tests for LGMD and related phenotypes.

## Acknowledgements

This publication was supported in part by the National Human Genome Research Institute of the National Institutes of Health through the following grant U24HG009650. The content is solely the responsibility of the author and does not necessarily represent the official views of the National Institutes of Health.

## References

1. Nigro V, Savarese M. Genetic basis of limb-girdle muscular dystrophies: the 2014 update. Acta Myol. 2014 May;33(1):1–12.

2. Strafella C, Campoli G, Galota RM, et al. Limb-Girdle Muscular Dystrophies (LGMDs): The Clinical Application of NGS Analysis, a Family Case Report. Front Neurol. 2019;10:619.

3. Angelini C. LGMD. Identification, description and classification. Acta Myol. 2020 Dec;39(4):207–17.

4. Georganopoulou DG, Moisiadis VG, Malik FA, et al. A Journey with LGMD: From Protein Abnormalities to Patient Impact. Protein J. 2021 Aug;40(4):466–88.

5. Straub V, Murphy A, Udd B, group Lws. 229th ENMC international workshop: Limb girdle muscular dystrophies Nomenclature and reformed classification Naarden, the Netherlands, 17-19 March 2017. Neuromuscul Disord. 2018 Aug;28(8):702–10.

6. Rehm HL, Berg JS, Brooks LD, et al. ClinGen--the Clinical Genome Resource. N Engl J Med. 2015 Jun 4;372(23):2235–42.

7. Strande NT, Riggs ER, Buchanan AH, et al. Evaluating the Clinical Validity of Gene-Disease Associations: An Evidence-Based Framework Developed by the Clinical Genome Resource. Am J Hum Genet. 2017 Jun 1;100(6):895–906.

8. Bean LJH, Funke B, Carlston CM, et al. Diagnostic gene sequencing panels: from design to report-a technical standard of the American College of Medical Genetics and Genomics (ACMG). Genet Med. 2020 Mar;22(3):453–61.

9. Thaxton C, Goldstein J, DiStefano M, et al. Lumping versus splitting: How to approach defining a disease to enable accurate genomic curation. Cell Genom. 2022 May 11;2(5).

10. Mungall CJ, McMurry JA, Kohler S, et al. The Monarch Initiative: an integrative data and analytic platform connecting phenotypes to genotypes across species. Nucleic Acids Res. 2017 Jan 4;45(D1):D712–D22.

11. Foley AR, Mohassel P, Donkervoort S, Bolduc V, Bonnemann CG. Collagen VI-Related Dystrophies. In: Adam MP, Feldman J, Mirzaa GM, Pagon RA, Wallace SE, Bean LJH, et al., editors. GeneReviews((R)). Seattle (WA)1993.

12. Vissing J, Barresi R, Witting N, et al. A heterozygous 21-bp deletion in CAPN3 causes dominantly inherited limb girdle muscular dystrophy. Brain. 2016 Aug;139(Pt 8):2154–63.

13. Raducu M, Baets J, Fano O, Van Coster R, Cruces J. Promoter alteration causes transcriptional repression of the POMGNT1 gene in limb-girdle muscular dystrophy type 2O. Eur J Hum Genet. 2012 Sep;20(9):945–52.

14. Clement EM, Godfrey C, Tan J, et al. Mild POMGnT1 mutations underlie a novel limb-girdle muscular dystrophy variant. Arch Neurol. 2008 Jan;65(1):137–41.

15. Song D, Dai Y, Chen X, et al. Genetic variations and clinical spectrum of dystroglycanopathy in a large cohort of Chinese patients. Clin Genet. 2021 Mar;99(3):384–95.

16. Dai Y, Liang S, Dong X, et al. Whole exome sequencing identified a novel DAG1 mutation in a patient with rare, mild and late age of onset muscular dystrophy-dystroglycanopathy. J Cell Mol Med. 2019 Feb;23(2):811–8.

17. Hara Y, Balci-Hayta B, Yoshida-Moriguchi T, et al. A dystroglycan mutation associated with limb-girdle muscular dystrophy. N Engl J Med. 2011 Mar 10;364(10):939–46.

18. Dong M, Noguchi S, Endo Y, et al. DAG1 mutations associated with asymptomatic hyperCKemia and hypoglycosylation of alpha-dystroglycan. Neurology. 2015 Jan 20;84(3):273–9.

19. Gundesli H, Talim B, Korkusuz P, et al. Mutation in exon 1f of PLEC, leading to disruption of plectin isoform 1f, causes autosomal-recessive limb-girdle muscular dystrophy. Am J Hum Genet. 2010 Dec 10;87(6):834–41.

20. Mroczek M, Durmus H, Topf A, Parman Y, Straub V. Four Individuals with a Homozygous Mutation in Exon 1f of the PLEC Gene and Associated Myasthenic Features. Genes (Basel). 2020 Jun 27;11(7).

21. Fattahi Z, Kahrizi K, Nafissi S, et al. Report of a patient with limb-girdle muscular dystrophy, ptosis and ophthalmoparesis caused by plectinopathy. Arch Iran Med. 2015 Jan;18(1):60–4.

22. Zhong J, Chen G, Dang Y, Liao H, Zhang J, Lan D. Novel compound heterozygous PLEC mutations lead to earlyITonset limbITgirdle muscular dystrophy 2Q. Mol Med Rep. 2017 May;15(5):2760–4.

23. Deev RV, Bardakov SN, Mavlikeev MO, et al. Glu20Ter Variant in PLEC 1f Isoform Causes Limb-Girdle Muscle Dystrophy with Lung Injury. Front Neurol. 2017;8:367.

24. Liang WC, Jong YJ, Wang CH, et al. Clinical, pathological, imaging, and genetic characterization in a Taiwanese cohort with limb-girdle muscular dystrophy. Orphanet J Rare Dis. 2020 Jun 23;15(1):160.

25. Vieira NM, Naslavsky MS, Licinio L, et al. A defect in the RNA-processing protein HNRPDL causes limb-girdle muscular dystrophy 1G (LGMD1G). Hum Mol Genet. 2014 Aug 1;23(15):4103–10.

26. Blazquez-Bernal A, Fernandez-Costa JM, Bargiela A, Artero R. Inhibition of autophagy rescues muscle atrophy in a LGMDD2 Drosophila model. FASEB J. 2021 Oct;35(10):e21914.

27. Bonnemann CG. The collagen VI-related myopathies Ullrich congenital muscular dystrophy and Bethlem myopathy. Handb Clin Neurol. 2011;101:81–96.

28. Nashabat M, Nabavizadeh N, Saracoglu HP, et al. SNUPN deficiency causes a recessive muscular dystrophy due to RNA mis-splicing and ECM dysregulation. Nat Commun. 2024 Feb 27;15(1):1758.

29. Iruzubieta P, Damborenea A, Ioghen M, et al. Biallelic variants in SNUPN cause a limb girdle muscular dystrophy with myofibrillar-like features. Brain. 2024 Feb 15.

30. Morales-Rosado JA, Schwab TL, Macklin-Mantia SK, et al. Bi-allelic variants in HMGCR cause an autosomal-recessive progressive limb-girdle muscular dystrophy. Am J Hum Genet. 2023 Jun 1;110(6):989–97.

31. Coppens S, Barnard AM, Puusepp S, et al. A form of muscular dystrophy associated with pathogenic variants in JAG2. Am J Hum Genet. 2021 May 6;108(5):840–56.

