## Supplementary material for "Expert Panel Curation of 31 Genes in Relation to Limb Girdle Muscular Dystrophy": Table1

| **HGNC gene symbol** | **OMIM entities (MIM#)** | **Old disease nomenclature** | **ENMC proposed nomenclature*** | **ClinGen curated disease entity (Mondo ID)** | **Classification (points)** | **Comments on precuration** | **LGMD GTR Panels^ including this gene** |
| --- | --- | --- | --- | --- | --- | --- | --- |
| *ANO5* | Miyoshi muscular dystrophy 3 (613319); Muscular dystrophy, limb-girdle, autosomal recessive 12 (611307); Gnathodiaphyseal dysplasia (166260) | LGMD 2L | LGMD R12 anoctamin5-related | autosomal recessive limb-girdle muscular dystrophy (MONDO_0015152) | Definitive  (12 GE, 6 EE, r/t) | Miyoshi muscular dystrophy and limb-girdle muscular dystrophy lumped, gnathodiaphyseal dysplasia split (may be curated by a different GCEP) | 22 |
| *BVES (POPDC1)* | Muscular dystrophy, limb-girdle, autosomal recessive 25 (616812) | LGMD 2X | BVES related myopathy | autosomal recessive limb-girdle muscular dystrophy (MONDO_0015152) | Definitive  (12 GE, 2 EE, r/t) |  | 5 |
| *CAPN3* | Muscular dystrophy, limb-girdle, autosomal dominant 4, Muscular dystrophy, limb-girdle, autosomal recessive 1 | LGMD 1I | LGMD D4 calpain3-related | autosomal dominant limb-girdle muscular dystrophy (MONDO:0015151) | Limited (1.4 GE, 1.75 EE) | Split curations for autosomal dominant and recessive entities | 22 |
| *CAPN3* | Muscular dystrophy, limb-girdle, autosomal dominant 4, Muscular dystrophy, limb-girdle, autosomal recessive 1 | LGMD 2A | LGMD R1 calpain3-related | autosomal recessive limb-girdle muscular dystrophy (MONDO:0015152) | Definitive  (12 GE, 6 EE, r/t) |  |  |
| *COL6A1* | Bethlem myopathy 1, Ullrich congenital muscular dystrophy 1 | Bethlem myopathy recessive | LGMD R22 collagen 6-related | collagen 6-related myopathy (MONDO:0100225), AR | Definitive  (12 GE, 6 EE, r/t) | Bethlem myopathy and Ullrich congenital muscular dystrophy lumped, but split curations for AR and AD inheritance patterns. Curated by Congenital Myopathies. | 3 |
| *COL6A1* | Bethlem myopathy 1, Ullrich congenital muscular dystrophy 1 | Bethlem myopathy dominant | LGMD D5 collagen 6-related | collagen 6-related myopathy (MONDO:0100225), AD | Definitive  (12 GE, 6 EE, r/t) |  |  |
| *COL6A2* | Bethlem myopathy 1, Ullrich congenital muscular dystrophy 1 | Bethlem myopathy recessive | LGMD R22 collagen 6-related | collagen 6-related myopathy (MONDO:0100225), AR | Definitive  (12 GE, 5 EE, r/t) | Bethlem myopathy and Ullrich congenital muscular dystrophy lumped, but split curations for AR and AD inheritance patterns. Curated by Congenital Myopathies. | 3 |
| *COL6A2* | Bethlem myopathy 1, Ullrich congenital muscular dystrophy 1 | Bethlem myopathy dominant | LGMD D5 collagen 6-related | collagen 6-related myopathy (MONDO:0100225), AD | Definitive  (12 GE, 3.75 EE, r/t) |  |  |
| *COL6A3* | Bethlem myopathy 1, Ullrich congenital muscular dystrophy 1 | Bethlem myopathy recessive | LGMD R22 collagen 6-related | collagen 6-related myopathy (MONDO:0100225), AR | Definitive  (12 GE, 3 EE, r/t) | Bethlem myopathy and Ullrich congenital muscular dystrophy lumped, but split curations for AR and AD inheritance patterns. Curated by Congenital Myopathies. | 3 |
| *COL6A3* | Bethlem myopathy 1, Ullrich congenital muscular dystrophy 1 | Bethlem myopathy dominant | LGMD D5 collagen 6-related | collagen 6-related myopathy (MONDO:0100225), AD | Definitive  (12 GE, 6 EE, r/t) |  |  |
| *DAG1* | Muscular dystrophy-dystroglycanopathy (congenital with brain and eye anomalies), type A, Muscular dystrophy-dystroglycanopathy (limb-girdle), type C, 9 | LGMD 2P | LGMD R16 α-dystroglycan-related | qualitative or quantitative defects of alpha-dystroglycan (MONDO:0018282), AR | Definitive (7 GE, 6 EE) | Both entities lumped into qualitative or quantitative defects of alpha-dystroglycan. Recuration in April, 2023, resulted in an upgrade from a Moderate to a Definitive classification. | 15 |
| *DNAJB6* | Muscular dystrophy, limb-girdle, autosomal dominant 1 (603511) | LGMD 1D | LGMD D1 DNAJB6-related | autosomal dominant limb-girdle muscular dystrophy (MONDO:0015151) | Definitive  (12 GE, 3 EE, r/t) |  | 21 |
| *DYSF* | Miyoshi muscular dystrophy 1 (254130); Muscular dystrophy, limb-girdle, autosomal recessive 2 (253601); Myopathy, distal, with anterior tibial onset (606768) | LGMD 2B | LGMD R2 dysferlin-related | autosomal recessive limb-girdle muscular dystrophy (MONDO:0015152) | Definitive  (12 GE, 6 EE, r/t) | All entities lumped into autosomal recessive limb-girdle muscular dystrophy | 22 |
| *FKRP* | Muscular dystrophy-dystroglycanopathy (congenital with brain and eye anomalies), type A, 5 (613153), Muscular dystrophy-dystroglycanopathy (congenital with or without impaired intellectual development), type B, 5 (606612), Muscular dystrophy-dystroglycanopathy (limb-girdle), type C, 5 (607155) | LGMD 2I | LGMD R9 FKRP-related | myopathy caused by variation in FKRP (MONDO:0700066), AR | Definitive  (12 GE, 6 EE, r/t) | All entities lumped into qualitative or quantitative defects of FKRP | 21 |
| *FKTN* | Cardiomyopathy, dilated, 1X, (611615); Muscular dystrophy-dystroglycanopathy (congenital with brain and eye anomalies), type A, 4 (253800); Muscular dystrophy-dystroglycanopathy (congenital without mental retardation), type B, 4 (613152); Muscular dystrophy-dystroglycanopathy (limb-girdle), type C, 4 (611588) | LGMD 2M | LGMD R13 Fukutin-related | myopathy caused by variation in FKTN (MONDO:0700067), AR | Definitive  (12 GE, 6 EE, r/t) | All entities lumped into myopathy caused by variation in FKTN | 21 |
| *GMPPB* | Muscular dystrophy-dystroglycanopathy (congenital with brain and eye anomalies), type A, 14 (615350), Muscular dystrophy-dystroglycanopathy (congenital with impaired intellectual development), type B, 14 (615351), Muscular dystrophy-dystroglycanopathy (limb-girdle), type C, 14 (615352) | LGMD 2T | LGMD R19 GMPPB-related | myopathy caused by variation in GMPPB (MONDO:0700084), AR | Definitive  (12 GE, 2.5 EE, r/t) | All entities lumped into myopathy caused by variation in GMPPB | 17 |
| *HNRNPDL* | Muscular dystrophy, limb-girdle, autosomal dominant 3 (609115) | LGMD 1G | LGMD D3 HNRNPDL-related | autosomal dominant limb-girdle muscular dystrophy (MONDO:0015151) | Moderate (7.1 GE, 1 EE) | Only one variant in HNRNPDL is observed in all individuals with AD-LGMD curated | 12 |
| *CRPPA (ISPD)* | Muscular dystrophy-dystroglycanopathy (congenital with brain and eye anomalies), type A, 7 (614643), Muscular dystrophy-dystroglycanopathy (limb-girdle), type C, 7 (616052) | LGMD 2U | LGMD R20 ISPD-related | myopathy caused by variation in CRPPA (MONDO:0100530), AR | Definitive  (12 GE, 5 EE, r/t) | All entities lumped into myopathy caused by variation in CRPPA | 15 |
| *LAMA2* | Muscular dystrophy, congenital, merosin deficient or partially deficient (607855), Muscular dystrophy, limb-girdle, autosomal recessive 23 (618138) | Laminin α2-related muscular dystrophy | LGMD R23 laminin α2-related | LAMA2-related muscular dystrophy (MONDO:0100228), AR | Definitive  (12 GE, 6 EE, r/t) | Both entities lumped into LAMA2-related muscular dystrophy. Curated by Congenital Myopathies and approved by LGMD GCEP. | 7 |
| *PLEC* | Epidermolysis bullosa simplex 5D, generalized intermediate, autosomal recessive (616487), Epidermolysis bullosa simplex 5A, Ogna type (131950), Epidermolysis bullosa simplex 5B, with muscular dystrophy (226670), Epidermolysis bullosa simplex 5C, with pyloric atresia (612138), Muscular dystrophy, limb-girdle, autosomal recessive 17 (613723) | LGMD 2Q | LGMD R17 plectin-related | autosomal recessive limb-girdle muscular dystrophy (MONDO:0015152) | Moderate (4.5 GE, 2 EE) | Split curation only for isolated LGMD phenotype due to biallelic variants impacting isoform 1f. Recuration in May 2023 resulted in no change in the Moderate classification. | 16 |
| *POGLUT1* | Dowling-Degos disease 4 (615696), Muscular dystrophy, limb-girdle, autosomal recessive 21 (617232) | LGMD 2Z | LGMD R21 POGLUT1-related | autosomal recessive limb-girdle muscular dystrophy (MONDO:0015152) | Definitive (10.8 GE, 1 EE, r/t) | Split curation only for autosomal recessive LGMD phenotype | 6 |
| *POMGNT1* | Muscular dystrophy-dystroglycanopathy (congenital with brain and eye anomalies), type A, 3 (253280), Muscular dystrophy-dystroglycanopathy (congenital with impaired intellectual development), type B, 3 (613151), Muscular dystrophy-dystroglycanopathy (limb-girdle), type C, 3 (613157), Retinitis pigmentosa 76 (617123) | LGMD 2O | LGMD R15 POMGnT1-related | myopathy caused by variation in POMGNT1 (MONDO:0700068), AR | Definitive (12 GE, 5.5 EE, r/t) | Retinitis pigmentosa split. Muscular dystrophy-dystroglycanopathy (congenital with brain and eye anomalies), type A, 3 and Muscular dystrophy-dystroglycanopathy (congenital with mental retardation), type B, 3 lumped into myopathy caused by variation in POMGNT1. Individuals reported in the literature with LGMD did not meet the LGMD GCEP-specific phenotype criteria and were not scored. | 17 |
| *POMGNT2* | Muscular dystrophy-dystroglycanopathy (congenital with brain and eye anomalies), type A, 8 (614830), Muscular dystrophy-dystroglycanopathy (limb-girdle) type C, 8 (618135) | POMGNT2-related muscular dystrophy | LGMD R24 POMGNT2-related | myopathy caused by variation in POMGNT2 (MONDO:0700069), AR | Definitive (8.5 GE, 6 EE, r/t) | All entities lumped into myopathy caused by variation in POMGNT2 | 7 |
| *POMT1* | Muscular dystrophy-dystroglycanopathy (congenital with brain and eye anomalies), type A, 1 (236670), Muscular dystrophy-dystroglycanopathy (congenital with impaired intellectual development), type B, 1 (613155), Muscular dystrophy-dystroglycanopathy (limb-girdle), type C, 1 (609308) | LGMD 2K | LGMD R11 POMT1-related | myopathy caused by variation in POMT1 (MONDO_0700070), AR | Definitive (12 GE, 4 EE, r/t) | All entities lumped into myopathy caused by variation in POMT1 | 21 |
| *POMT2* | autosomal recessive congenital muscular dystrophy-dystroglycanopathy with brain and eye anomalies type A2 (613150); autosomal recessive congenital muscular dystrophy-dystroglycanopathy with mental retardation type B2 (613156); autosomal recessive limb-girdle muscular dystrophy-dystroglycanopathy type C2 (613158) | LGMD 2N | LGMD R14 POMT2-related | myopathy caused by varation in POMT2 (MONDO_0700071), AR | Definitive (12 GE, 5.5 EE, r/t) | All entities lumped into myopathy caused by variation in POMT2 | 18 |
| *POPDC3* | Muscular dystrophy, limb-girdle, autosomal recessive 26 (618848) | none | none | autosomal recessive limb-girdle muscular dystrophy (MONDO_0015152) | Moderate (6.8 GE, 1 EE, r/t) | Recuration in April, 2023, resulted in change Limited to Moderate classification | 0 |
| *SGCA* | Muscular dystrophy, limb-girdle, autosomal recessive 3 (608099) | LGMD 2D | LGMD R3 α-sarcoglycan-related | autosomal recessive limb-girdle muscular dystrophy (MONDO_0015152) | Definitive (12 GE, 6 EE, r/t) |  | 22 |
| *SGCB* | Muscular dystrophy, limb-girdle, autosomal recessive 4 (604286) | LGMD 2E | LGMD R4 β-sarcoglycan-related | autosomal recessive limb-girdle muscular dystrophy (MONDO_0015152) | Definitive (12 GE, 5.5 EE, r/t) |  | 22 |
| *SGCD* | Cardiomyopathy, dilated, 1L (606685), Muscular dystrophy, limb-girdle, autosomal recessive 6 (601287) | LGMD 2F | LGMD R6 δ-sarcoglycan-related | autosomal recessive limb-girdle muscular dystrophy (MONDO_0015152) | Definitive (12 GE, 5 EE, r/t) | Dilated cardiomyopathy lumped with autosomal recessive limb-girdle muscular dystrophy | 22 |
| *SGCG* | Muscular dystrophy, limb-girdle, autosomal recessive 5 (253700) | LGMD 2C | LGMD R5 γ-sarcoglycan-related | autosomal recessive limb-girdle muscular dystrophy (MONDO_0015152) | Definitive (12 GE, 6 EE, r/t) |  | 22 |
| *TCAP* | Cardiomyopathy, hypertrophic, 25 607487 AD 3 Muscular dystrophy, limb-girdle, autosomal recessive 7 601954 | LGMD 2G | LGMD R7 telethonin-related | autosomal recessive limb-girdle muscular dystrophy (MONDO_0015152) | Definitive (12 GE, 3.5 EE, r/t) | Hypertrophic cardiomyopathy split from autosomal recessive limb-girdle muscular dystrophy | 22 |
| *TNPO3* | Muscular dystrophy, limb-girdle, autosomal dominant 2 (608423) | LGMD 1F | LGMD D2 TNP03-related | autosomal dominant limb-girdle muscular dystrophy (MONDO_0015151) | Moderate (4.5 GE, 2 EE) | Recuration in April, 2023, resulted in an upgrade from a Limited to a Moderate classification. | 14 |
| *TRAPPC11* | Muscular dystrophy, limb-girdle, autosomal recessive 18 (615356) | LGMD 2S | LGMD R18 TRAPPC11-related | autosomal recessive limb-girdle muscular dystrophy (MONDO_0015152) | Definitive (12 GE, 2.5 EE, r/t) |  | 17 |
| *TRIM32* | ?Bardet-Biedl syndrome 11 (615988), Muscular dystrophy, limb-girdle, autosomal recessive 8 (254110) | LGMD 2H | LGMD R8 TRIM 32-related | autosomal recessive limb-girdle muscular dystrophy (MONDO_0015152) | Definitive (9.6 GE, 4 EE, r/t) | Split Bardet-Biedl syndrome 11. Curated only for autosomal recessive limb-girdle muscular dystrophy | 22 |
| *TTN* | Cardiomyopathy, dilated, 1G (604145), Cardiomyopathy, familial hypertrophic, 9 (613765), Muscular dystrophy, limb-girdle, autosomal recessive 10 (608807), Myopathy, myofibrillar, 9, with early respiratory failure (603689), Salih myopathy (611705), Tibial muscular dystrophy, tardive (600334) | LGMD 2J | LGMD R10 titin-related | TTN-related myopathy (MONDO:0100175), AR | Definitive (12 GE, 6 EE, r/t) | Recessive forms of titinopathies lumped into TTN-related myopathy: limb-girdle muscular dystrophy, centronuclear myopathy, Salih myopathy (MIM #611705), Emery-Dreifuss-like muscular dystrophy, titinopathy with congenital contractures, minicore myopathy, and distal titinopathy. Curated by Congenital Myopathies GCEP | 20 |
| **229^th^ ENMC International workshop (Straub et al, 2018; PMID: 30055862)* | | | | | | | |
| *^A total of 22 GTR panels were identified that incldued 'limb girdle muscular dystrophy', 'limb-girdle muscular dystrophy', 'LGMD' in the panel name in the NIH genetic testing registry (GTR), queried on 10/25/2023. Panel names indicating AR (5) and AD (3) were also identified, but not included in the denominator.* | | | | | | | |
